# Involvement of the splicing factor SART1 in the BRCA1-dependent homologous recombination repair of DNA double-strand breaks

**DOI:** 10.1101/2023.09.07.556638

**Authors:** Motohiro Yamauchi, Reona Kato, Takaaki Yasuhara, Yuki Uchihara, Miyako Hirakawa, Kie Ozaki, Yu Abe, Hiroki Shibata, Reika Kawabata-Iwakawa, Aizhan Shakayeva, Palina Kot, Kiyoshi Miyagawa, Keiji Suzuki, Atsushi Shibata, Naoki Matsuda

## Abstract

Although previous studies have reported that pre-mRNA splicing factors (SFs) are involved in the repair of DNA double-strand breaks (DSBs) via homologous recombination (HR), their exact role in promoting HR remains poorly understood. Here, we showed that SART1, an SF upregulated in several types of cancer, promotes DSB end resection, which is an essential step in HR, without affecting the expression of HR factors. In the promotion of resection, SART1 is epistatic with BRCA1, an HR factor known to promote resection. Mechanistically, SART1 cooperates with BRCA1 to counteract the resection blockade posed by 53BP1 and RIF1. Moreover, SART1 is required for recruitment of the HR-promoting portion of BRCA1 to DSBs. Laser microirradiation experiments revealed that SART1 is recruited to DSBs in a BRCA1-dependent manner, and that the arginine/serine (RS)-rich domain is required for SART1 recruitment to DSBs. We also obtained evidence that SART1 is involved in transcription-associated HR together with BRCA1. Furthermore, chromosome analysis demonstrated that SART1 and BRCA1 epistatically suppressed genomic alterations caused by DSB misrepair in the G2 phase. Collectively, these results indicate that SART1 and BRCA1 cooperatively facilitate resection of DSBs arising in transcriptionally active genomic regions, thereby promoting faithful HR repair of such DSBs and suppressing genome instability.

## Introduction

Dysregulated RNA splicing has recently been linked to both the promotion and suppression of tumorigenesis, as mutations and altered expression (both upregulation and downregulation) of splicing factors (SFs) have been found in a variety of human malignancies (Bradley and Anczuków, 2023). In addition to their central role in shaping the transcriptome, recent studies have revealed the involvement of SFs in suppressing genome instability, which is a hallmark of cancer (Hanahan, 2022). For example, a previous study demonstrated that small interfering RNA (siRNA)-mediated knockdown of SFs leads to elevated levels of histone H2AX phosphorylation at serine 139, which represents DNA damage, including DNA double-strand breaks (DSBs) (Paulsen et al., 2009). Moreover, two genome-wide screening studies have identified many SFs as factors promoting homologous recombination (HR), which is a major DSB repair pathway (Adamson et al., 2012; Herr et al., 2015). Previous studies have shown that both the splicing-dependent and splicing-independent functions of SFs contribute to HR promotion (Anantha et al., 2013; Awwad et al., 2023; MacHour et al., 2021; Onyango et al., 2017; Prados-Carvajal et al., 2018; Tanikawa et al., 2016). However, the splicing-independent function (i.e., the direct function) of SFs in HR has not been fully studied.

In addition to HR, there is another major DSB repair pathway, non-homologous end-joining (NHEJ). NHEJ is active throughout the cell cycle, whereas HR operates mainly in the S/G2 phase when sister chromatids are available (Hustedt and Durocher, 2017). In somatic cells, HR is believed to be accurate because conservative synthesis-dependent strand annealing, an HR sub-pathway, is predominant in this cell type (Scully et al., 2019). Thus, in the S/G2 phase, both NHEJ and HR can operate, and the mechanism of choice between these two pathways has been extensively studied. Accumulating evidence indicates that DSB end resection, which generates the 3’-overhang required to initiate HR, is a critical determinant of the pathway choice between NHEJ and HR (Ceccaldi et al., 2016; Scully et al., 2019; Shibata, 2017). It is known that BRCA1 directs the repair pathway toward HR by promoting resection (Tarsounas and Sung, 2020). BRCA1 antagonizes 53BP1, which inhibits resection (Panier and Boulton, 2014). Upon DSBs, 53BP1 is phosphorylated by ATM kinase, and phosphorylated 53BP1 recruits RIF1, a downstream effector of 53BP1, to DSBs, thereby blocking resection (Panier and Boulton, 2014). BRCA1 facilitates resection by promoting protein phosphatase 4C-dependent 53BP1 dephosphorylation and RIF1 release from DSBs (Isono et al., 2017). However, the involvement of SFs in the antagonism between BRCA1 and 53BP1 over the choice of repair pathway remains largely unknown.

Among the SFs implicated in HR in previous studies, we focused on SART1 in this study because of its upregulated expression in several types of human cancer and its correlation with tumor resistance to DNA-damaging agents (Adamson et al., 2012; Allen et al., 2012; Kawamoto et al., 1999; Sasatomi et al., 2000; Shintaku et al., 2000). In this study, we revealed that SART1 is recruited to DSB sites and promotes resection of DSBs, particularly those arising in transcriptionally active genomic regions. This resection-promoting function of SART1 appears to be closely interlinked with that of BRCA1, where SART1 operates epistatically with BRCA1 to counteract 53BP1/RIF1-mediated resection blockade. Moreover, our results demonstrate that SART1 is required for recruiting the HR-promoting portion of BRCA1 to DSBs. Consistent with the epistatic relationship between SART1 and BRCA1 during resection, these factors cooperatively suppress genomic alterations caused by DSB misrepair in the G2 phase. These results indicate that when DSBs occur in transcriptionally active genomic regions, SART1 is rapidly recruited to DSB sites and recruits BRCA1, which facilitates resection by antagonizing the 53BP1/RIF1 resection barrier, thereby promoting faithful DSB repair by HR.

## Results

### SART1 promotes HR by facilitating DSB end resection

This study aimed to investigate the mechanism by which SART1 promotes HR. First, we reassessed the involvement of SART1 in HR using a well-established HR reporter assay (direct repeat (DR)-green fluorescent protein (GFP) assay) (Pierce et al., 1999). This assay uses a human osteosarcoma U2OS cell line with a chromosomally integrated recognition site for the I-Sce I endonuclease, and the cells express GFP when the I-Sce I-induced DSB is repaired by HR. As shown in Figs. 1a–b, siRNA-mediated knockdown of SART1 significantly decreased the percentage of GFP(+) cells without affecting cell cycle distribution, confirming the previous finding that SART1 is involved in HR (Adamson et al., 2012). Next, RNA sequencing (RNA-seq) was performed to determine the effect of SART1 knockdown on the transcriptome of human telomerase reverse transcriptase-immortalized retinal pigment epithelial (RPE-hTERT) cells. Analysis of RNA-seq data demonstrated that SART1 knockdown did not affect the mRNA expression levels of most genes, including HR factors (Figs. S1a–b). Furthermore, SART1 knockdown did not affect the protein levels of major HR factors, such as MRE11, CtIP, BRCA1, and RAD51, in RPE-hTERT cells (Fig. S2). To further confirm the involvement of SART1 in HR, we examined nuclear foci of RAD51, another established indicator of HR (Herr et al., 2015). RAD51 foci in the G2 phase cells after exposure to ionizing radiation (IR) were analyzed because in this study, we mostly focused on HR of two-ended DSBs induced by IR in the G2 phase. To identify G2 phase cells, CENPF protein (S/G2 phase marker) and 5-ethynyl-2′-deoxyuridine (EdU; S phase marker) were concomitantly visualized and EdU(−)/CENPF(+) cells were regarded as G2 phase cells (Fig. 1c). SART1 knockdown significantly decreased the number of IR-induced RAD51 foci (Figs. 1c–d and S3a). Next, we examined the role of SART1 in DSB end resection, an essential first step in HR (Cejka and Symington, 2021). To visualize the sites of DSBs that underwent resection, the nuclear foci of replication protein A (RPA), a widely used indicator of resected DSBs, was employed (Isono et al., 2017; Yasuhara et al., 2018). SART1 knockdown decreased the number of RPA foci in G2-irradiated cells (Figs. 1e–f and S3b). Collectively, these results indicate that SART1 promotes HR by facilitating DSB end resection.

**Figure 1.**
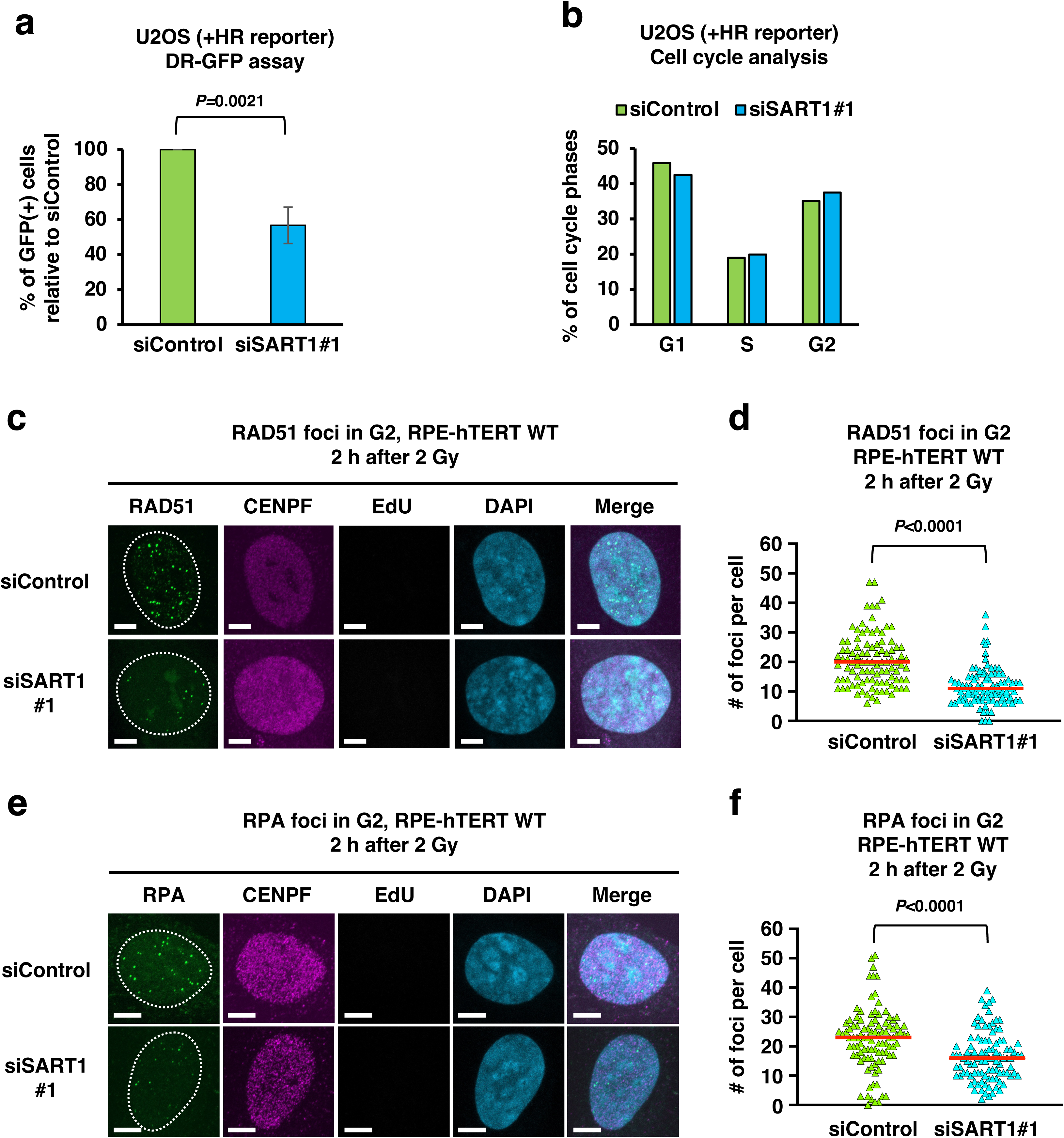
SART1 promotes homologous recombination (HR) by facilitating DSB end resection. (a) Effect of SART1 knockdown on HR efficiency. U2OS cells harboring the HR reporter were transfected with the indicated siRNA and subjected to the DR-GFP assay. The percentage of siSART1#1-transfected GFP(+) cells relative to that of siControl-transfected GFP(+) cells is shown as mean ± standard deviation (N = 3). (b) Effect of SART1 knockdown on cell cycle. U2OS cells harboring the HR reporter were transfected with the indicated siRNA. Cells were then fixed, and the nuclei were stained with propidium iodide. The cell cycle status was identified by flow cytometry. (c) Immunofluorescence images of RAD51 foci in SART1 knockdown cells. Wild-type RPE-hTERT cells (RPE-hTERT WT) were transfected with indicated siRNAs. Two days after siRNA transfection, cells were irradiated with 2 Gy γ-rays and fixed 2 h later. The cells were treated with EdU 30 m before irradiation until fixation to label S-phase cells. Fixed cells were subjected to RAD51/CENPF immunofluorescence and EdU detection. RAD51 foci in CENPF(+)/EdU(−) G2 phase cells are shown. (d) Number of RAD51 foci in SART1 knockdown cells. Immunofluorescence samples were prepared as described in (c). (e) Immunofluorescence images of RPA foci in SART1 knockdown cells. siRNA transfection, EdU treatment, irradiation, and fixation were performed, as described in (c). Fixed cells were subjected to RPA/CENPF immunofluorescence and EdU detection. RPA foci in CENPF(+)/EdU(−) G2 phase cells are shown. (f) Number of RPA foci in SART1 knockdown cells. Immunofluorescence samples were prepared as described in (e). In (d) and (f), the number of foci in 100 G2 cells from two independent experiments (50 G2 cells/experiment/sample) is shown. Each symbol in (d) and (f) represents the number of foci per cell. Red bars in (d) and (f) indicate the median number of foci in each sample. Scale bars in (c) and (e):5 µm.

### SART1 promotes resection in an epistatic manner with BRCA1

It is well-established that BRCA1 promotes resection (Tarsounas and Sung, 2020). Therefore, we next examined the role of SART1 in BRCA1-dependent resection. Single knockdown of BRCA1 and SART1 similarly reduced the number of IR-induced RPA foci in the G2 phase (Fig. 2a). The double knockdown of BRCA1 and SART1 did not additively decrease the number of RPA foci when compared to the single knockdown of BRCA1 or SART1, indicating that these two proteins epistatically promote resection (Fig. 2a). Consistently, the number of RAD51 foci was similarly reduced by the single knockdown of BRCA1 and SART1, and the double knockdown of these proteins did not show an additive effect, indicating epistasis between BRCA1 and SART1 in HR (Fig. 2b).

**Figure 2.**
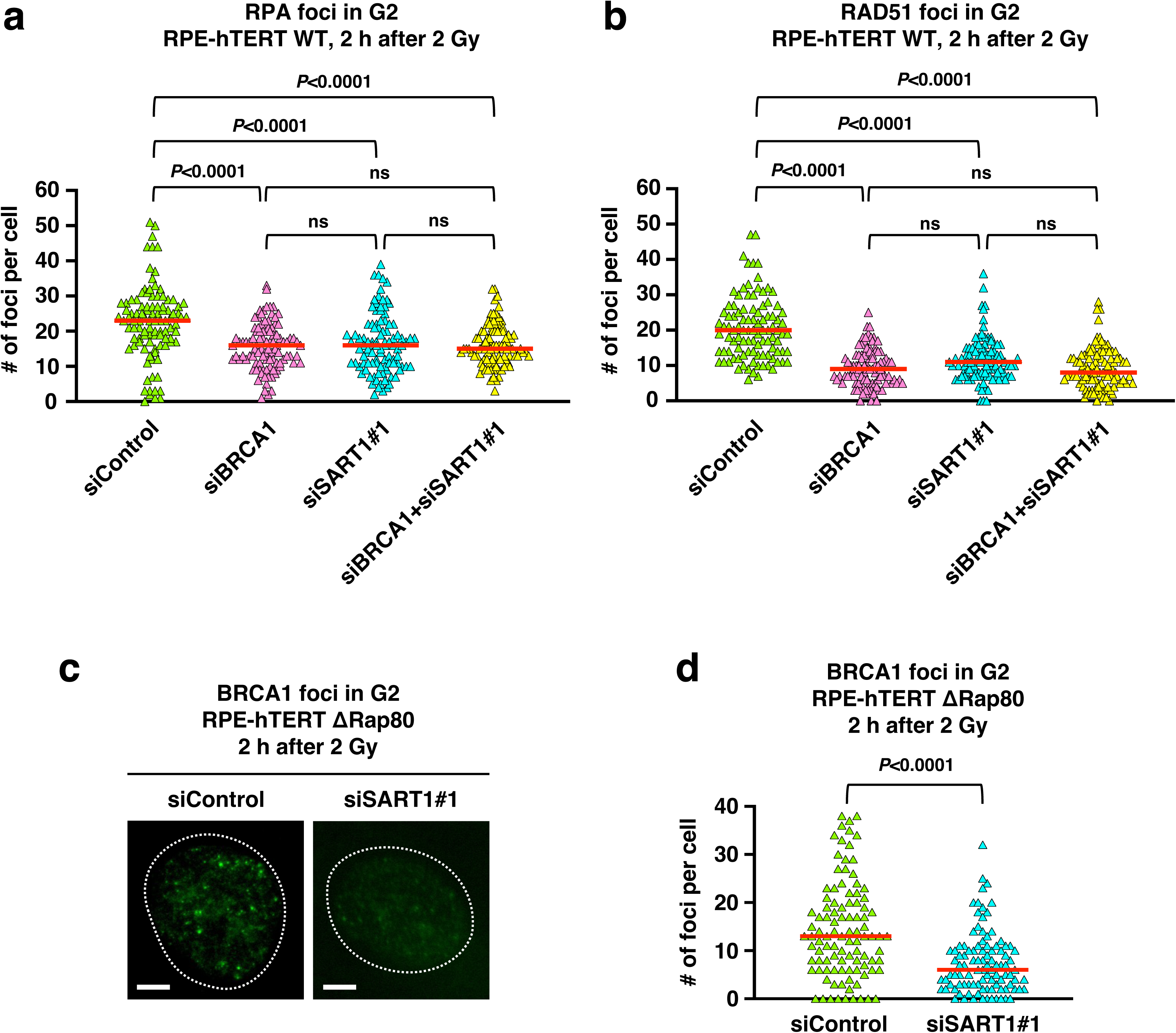
SART1 promotes resection in an epistatic manner with BRCA1. (a) Number of RPA foci in BRCA1 and/or SART1 knockdown cells. RPE-hTERT WT cells were transfected with indicated siRNAs (s). Two days after siRNA transfection, the cells were irradiated with 2 Gy γ-rays and fixed 2 h later. The cells were treated with EdU from 30 m before irradiation until fixation to label S phase cells. The fixed cells were subjected to RPA/CENPF immunofluorescence and EdU detection. The number of RPA foci in the CENPF(+)/EdU(−) G2 phase cells was counted. (b) Number of RAD51 foci in BRCA1 and/or SART1 knockdown cells. siRNA transfection, EdU treatment, irradiation, and fixation were performed as described in (a). The fixed cells were subjected to RAD51/CENPF immunofluorescence and EdU detection. The number of RAD51 foci in the CENPF(+)/EdU(−) G2 phase cells was counted. (c) Immunofluorescence images of BRCA1 foci in Rap80-knockout cells. Rap80-knockout RPE-hTERT cells (RPE-hTERT ΔRap80) were transfected with indicated siRNAs. Two days after siRNA transfection, the cells were irradiated with 2 Gy γ-rays and fixed 2 h later. The cells were treated with EdU from 30 m before irradiation until fixation to label S phase cells. Fixed cells were subjected to BRCA1/CENPF immunofluorescence and EdU detection. BRCA1 foci in CENPF(+)/EdU(−) G2 phase cells are shown. (d) Effect of SART1 knockdown on the number of BRCA1 foci in Rap80-knockout cells. The immunofluorescence samples were prepared as described in (c). The number of BRCA1 foci in the CENPF(+)/EdU(−) G2 phase cells was counted. In (a), (b), and (d), the number of foci in 100 G2 cells from two independent experiments (50 G2 cells/experiment/sample) is shown. Each symbol in (a), (b) and (d) represents the number of foci per cell. Red bars in (a), (b) and (d) indicate the median number of foci in each sample. Scale bars in (c):5 µm.

### SART1 promotes the recruitment of the HR-promoting portion of BRCA1 to DSBs

Upon DSBs, BRCA1 is recruited to DSB sites and forms discrete nuclear foci (Hu et al., 2011). To gain more insight into the role of SART1 in BRCA1-dependent resection, we examined the impact of SART1 knockdown on the recruitment of BRCA1 to DSBs by assessing BRCA1 foci formation. At DSB sites, BRCA1 forms several distinct protein complexes; among them, the BRCA1-Rap80 complex suppresses HR, whereas other complexes promote HR (Li and Greenberg, 2012). To visualize recruitment of only HR-promoting BRCA1, the number of BRCA1 foci in cells lacking Rap80 (RPE-hTERT ΔRap80 cells) was examined. As shown in Figs. 2c–d, SART1 knockdown significantly reduced the number of BRCA1 foci in G2-irradiated ΔRap80 cells, indicating that SART1 promotes the recruitment of the HR-promoting portion of BRCA1 to DSBs.

### SART1 and BRCA1 epistatically counteract 53BP1/RIF1-mediated resection blockade

Current evidence indicates that BRCA1 promotes resection by antagonizing the resection barrier posed by 53BP1 and its downstream effectors such as RIF1 (Scully et al., 2019; Shibata, 2017; Tarsounas and Sung, 2020). Thus, we next examined the role of SART1 in this BRCA1’s function. A previous study reported that loss of 53BP1 restores impaired resection and RAD51 foci in BRCA1-deficient cells (Bunting et al., 2010). Consistent with this, 53BP1 knockdown restored the reduced number of RPA foci in BRCA1 knockdown cells (Fig. 3a, cf. siBRCA1 and siBRCA1+si53BP1). Similarly, 53BP1 knockdown reversed the reduction in the number of RPA foci in SART1 knockdown cells (Fig. 3a, cf. siSART1#1 and siSART1#1+si53BP1). The knockdown of 53BP1 also restored the number of RAD51 foci in BRCA1 and SART1 knockdown cells (Fig. 3b). A previous study demonstrated that BRCA1 promotes 53BP1 dephosphorylation and RIF1 release from DSBs (Isono et al., 2017). Thus, we next examined the role of SART1 in BRCA1-dependent 53BP1 dephosphorylation and RIF1 release from DSBs. Nuclear foci of threonine 543-phosphorylated 53BP1 (53BP1 pT543) were used to monitor 53BP1 phosphorylation at DSB sites. At both early (30 m) and late (4 h) time points after IR, the number of 53BP1 pT543 foci was significantly higher in BRCA1- or SART1 single knockdown cells or BRCA1/SART1 double knockdown cells than in control cells (Figs. 3c–d). The number of 53BP1 pT543 foci was comparable between BRCA1 knockdown and SART1 knockdown cells, and the BRCA1/SART1 double knockdown did not additively increase the number of 53BP1 pT543 foci compared with the single knockdown of BRCA1 or SART1, at both 30 m and 4 h after IR (Fig. 3c).

**Figure 3.**
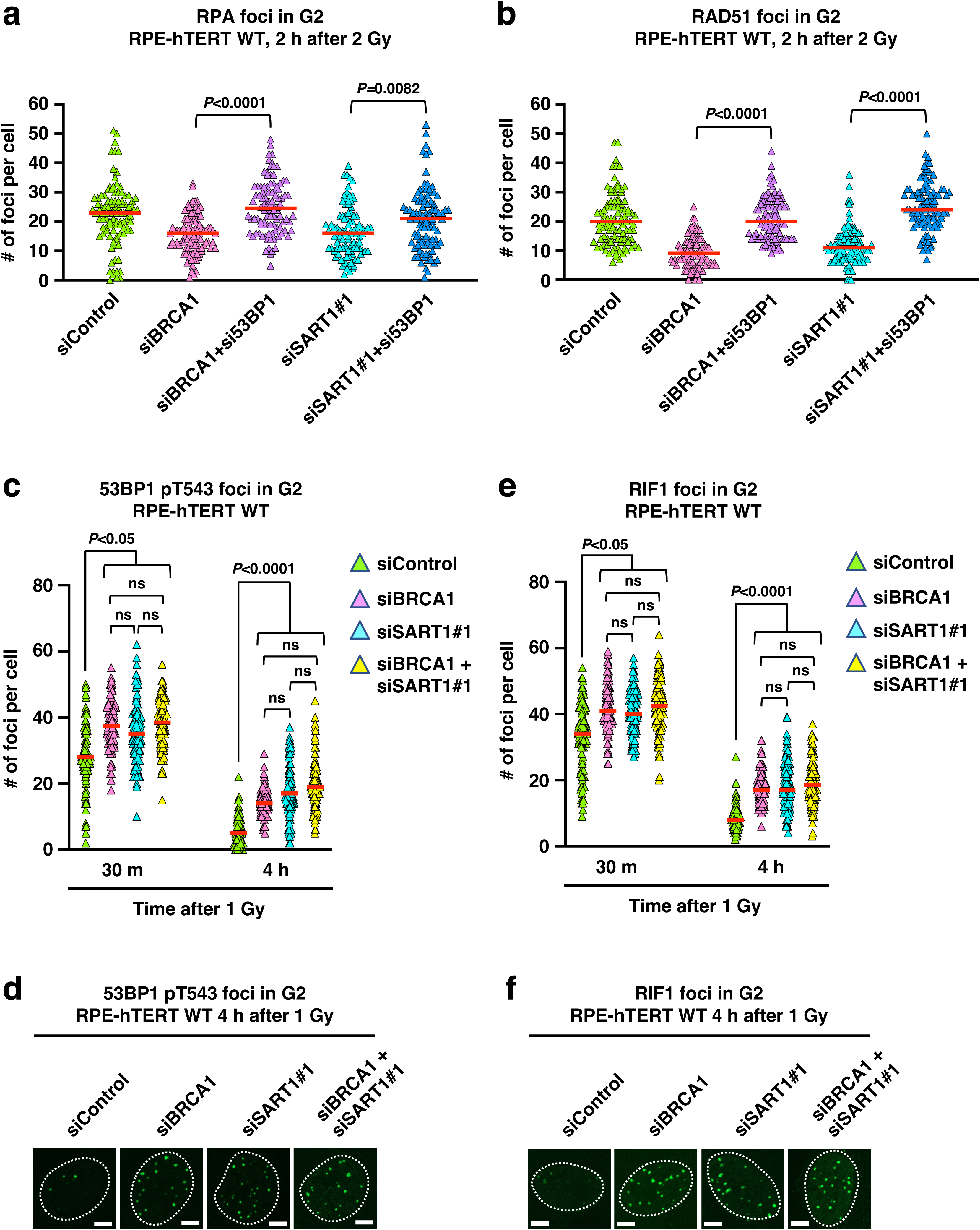
SART1 and BRCA1 epistatically counteract the 53BP1/RIF1-mediated resection blockade. (a) Effect of 53BP1 knockdown on the number of RPA foci in BRCA1 or SART1 knockdown cells. RPE-hTERT WT cells were transfected with the indicated siRNA(s). Two days after siRNA transfection, the cells were irradiated with 2 Gy γ-rays and fixed 2 h later. The cells were treated with EdU from 30 m before irradiation until fixation to label S phase cells. The fixed cells were subjected to RPA/CENPF immunofluorescence and EdU detection. The number of RPA foci in CENPF(+)/EdU(−) G2 phase cells was counted. (b) Effect of 53BP1 knockdown on the number of RAD51 foci in BRCA1 or SART1 knockdown cells. siRNA transfection, EdU treatment, irradiation, and fixation were performed as described in (a). The fixed cells were subjected to RAD51/CENPF immunofluorescence and EdU detection. The number of RAD51 foci in CENPF(+)/EdU(−) G2 phase cells was counted. (c) Effect of BRCA1 and/or SART1 knockdown on 53BP1 phosphorylation at DSBs. RPE-hTERT WT cells were transfected with the indicated siRNA(s). Two days after siRNA transfection, the cells were irradiated with 1 Gy γ-rays and fixed 30 m or 4 h later. The cells were treated with EdU from 30 m before irradiation until fixation to label S phase cells. Fixed cells were subjected to immunofluorescence of phosphorylated 53BP1 at threonine 543 (53BP1 pT543) and CENPF followed by EdU detection. The number of 53BP1 pT543 foci in CENPF(+)/EdU(−) G2 phase cells was counted. (d) Immunofluorescence images of 53BP1 pT543 foci 4 h after irradiation. Samples were prepared as described in (c). 53BP1 pT543 foci in CENPF(+)/EdU(−) G2 phase cells are shown. (e) Effect of BRCA1 and/or SART1 knockdown on RIF1 foci formation at DSBs. The siRNA transfection, EdU treatment, irradiation and fixation were done as described in (c). Fixed cells were subjected to RIF1/CENPF immunofluorescence and EdU detection. The number of RIF1 foci in the CENPF(+)/EdU(−) G2 phase cells was counted. (f) Immunofluorescence images of RIF1 foci 4 h after irradiation. The samples were prepared as described in (e). RIF1 foci in CENPF(+)/EdU(−) G2 phase cells are shown. In (a)–(c) and (e), the number of foci in 100 G2 cells from two independent experiments (50 G2 cells/experiment/sample) is shown. Each symbol in (a)-(c) and (e) represents the number of foci per cell. Red bars in (a)–(c) and (e) indicate the median number of foci in each sample. Scale bars in (d) and (f): 5 µm.

Next, we examined the impact of BRCA1 and/or SART1 knockdown on RIF1 foci, an indicator of RIF1 localization at DSBs (Escribano-Díaz et al., 2013; Isono et al., 2017). Similar to the number of 53BP1 pT543 foci, the number of RIF1 foci in BRCA1 or SART1 single knockdown cells was significantly higher than that in control cells at both 30 m and 4 h after IR (Figs. 3e–f). BRCA1/SART1 double knockdown did not further increase the number of RIF1 foci compared to BRCA1 or SART1 single knockdown at either time point (Fig. 3e). These results indicate that SART1 and BRCA1 epistatically counteract the 53BP1/RIF1-mediated resection blockade by facilitating 53BP1 dephosphorylation and RIF1 release from DSBs.

### SART1 is recruited to DSB sites in a BRCA1-dependent manner

If SART1 plays a direct role in HR, it should be recruited to DSBs. Therefore, we performed live-cell imaging to examine SART1 recruitment to DSBs using a near-infrared (730 nm) two-photon microbeam that induces DSBs in the presence of a photosensitizer (Reynolds et al., 2013; Yasuhara et al., 2018). When U2OS cells expressing GFP-SART1 and mCherry-Geminin (a marker of S/G2 phases) were irradiated with the laser, GFP-SART1 accumulated along the laser track in S/G2 phase cells (Fig. 4a). Accumulation began 10 s after irradiation, peaked at 30–40 s, and persisted for at least 110 s (Fig. 4b). Consistent with the epistatic relationship between SART1 and BRCA1 in resection and HR, BRCA1 knockdown significantly decreased the recruitment of SART1 along the laser track (Fig. 4c).

**Figure 4.**
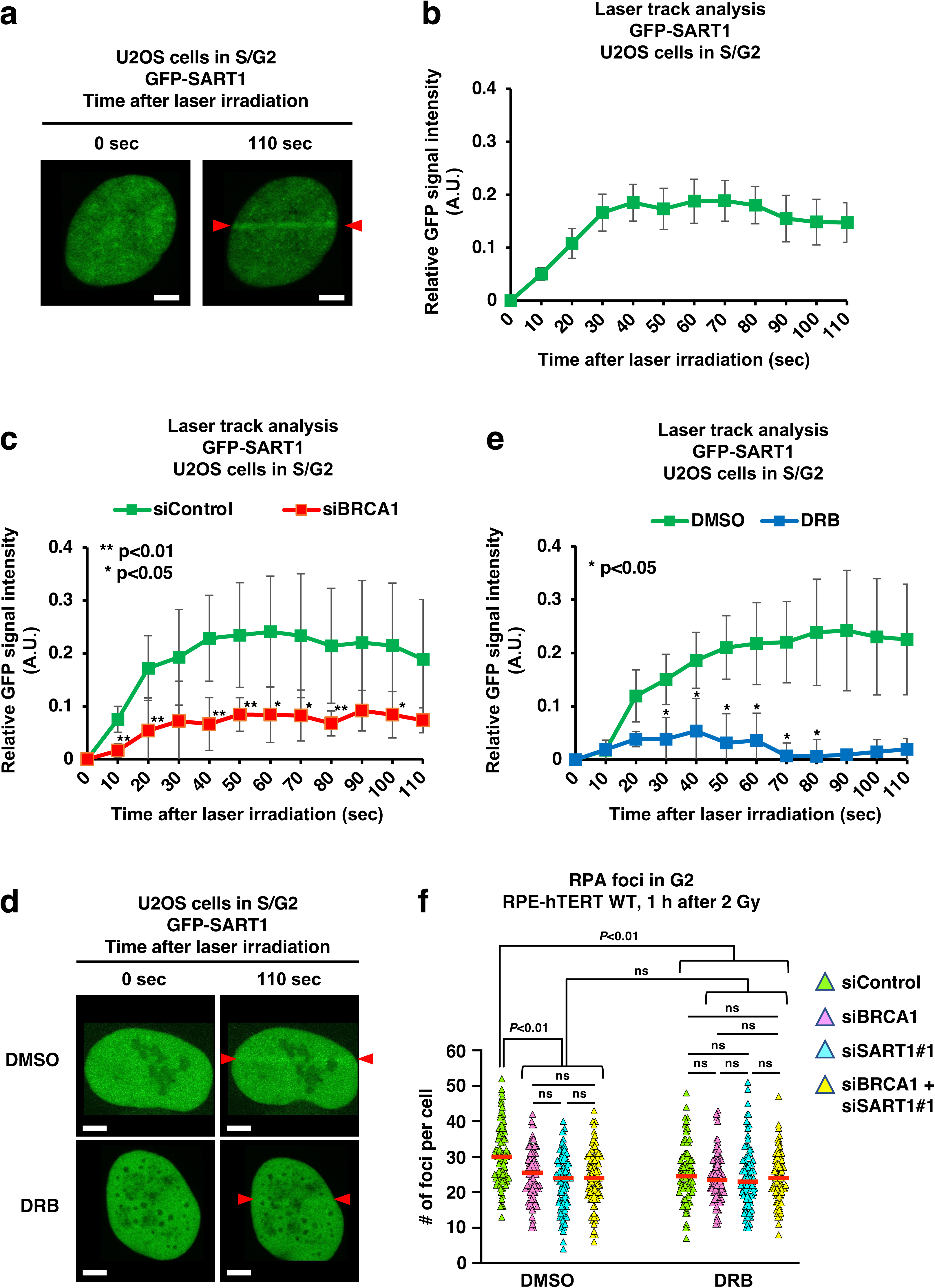
SART1 is recruited to DSB sites and promotes transcription-associated HR in an epistatic manner with BRCA1. (a) SART1 is recruited to laser-induced DSBs. U2OS cells expressing mCherry-Geminin (an S/G2 phase marker) were transfected with GFP-SART1 vector. One day later, GFP(+)/mCherry(+) cells were irradiated with the 730 nm laser along the line flanked by two red arrowheads. A photosensitizer (Hoechst33342, 10 µg/mL) was added 30 m before irradiation. (b) Kinetics of SART1 recruitment to the laser tracks. Vector transfection and laser irradiation were performed as described in (a). The intensity of GFP-SART1 in the laser-irradiated regions of S/G2-phase U2OS cells was recorded every 10 s until 110 s after irradiation. (c) SART1 recruitment to the laser tracks is BRCA1-dependent. U2OS cells expressing mCherry-Geminin were transfected with indicated siRNAs. Two days after siRNA transfection, the cells were transfected and irradiated as described in (a). The intensity of GFP-SART1 was recorded as described in (b). (d) Effect of transcription inhibition on SART1 recruitment to laser tracks. Vector transfection and laser irradiation were done as described in (a). The cells were treated with DRB (100 µM) or DMSO (vehicle control) from 30 m before irradiation. (e) Transcription inhibition suppresses SART1 recruitment to the laser tracks. Vector transfection, chemical treatment, and laser irradiation were performed, as described in (d). The intensity of GFP-SART1 was recorded as descried in (b). (f) SART1 promotes transcription-associated resection in an epistatic manner with BRCA1. RPE-hTERT WT cells were transfected with the indicated siRNA(s). Two days after siRNA transfection, cells were irradiated with 2 Gy γ-rays and fixed 1 h later. DRB (100 µM) or DMSO was added 30 m before irradiation until fixation. EdU was applied 30 m before irradiation until fixation to label S phase cells. The fixed cells were subjected to RPA/CENPF immunofluorescence and EdU detection. The number of RPA foci in the CENPF(+)/EdU(−) G2 phase cells is shown. Scale bars in (a) and (d): 5 µm. Error bars in (b), (c), and (e) represent standard deviation (SD); N = 3−6. *, p < 0.05. Red bars in (f) indicate the median number of foci in each sample.

### SART1 promotes transcription-associated HR in an epistatic manner with BRCA1

We previously discovered an HR sub-pathway that repairs DSBs arising in transcriptionally active genomic regions in the G2 phase and found that BRCA1 is involved in this transcription-associated HR (Yasuhara et al., 2018). Considering that SFs, such as SART1, should be located at transcription sites, we hypothesized that SART1 also participates in transcription-associated HR. We first examined whether the transcription status affected the recruitment of SART1 to DSB sites. Laser irradiation experiments revealed that SART1 recruitment to the laser track was significantly decreased in the presence of 5,6-dichloro-1-b-D-ribofuranosylbenzimidazole (DRB), an inhibitor of transcription elongation by RNA polymerase II (Figs. 4d–e) (Yamaguchi et al., 1999). We previously demonstrated that transcriptional inhibition by DRB reduced the number of RPA foci in G2-irradiated cells, indicating that active transcription is required for the resection of a subset of DSBs (Yasuhara et al., 2018). Therefore, we examined the role of SART1 in transcription-dependent resection. Consistent with our previous results, transcriptional inhibition by DRB reduced the number of RPA foci in siControl-transfected cells (Fig. 4f, cf. siControl DMSO and siControl DRB). However, the single knockdown of BRCA1 or SART1 or the double knockdown of BRCA1/SART1 did not further decrease the number of RPA foci in DRB-treated cells (Fig. 4f, cf. siControl, siBRCA1, siSART1#1, and siBRCA1+siSART1#1 in the DRB-treated group). Moreover, DRB treatment did not additively decrease RPA foci in BRCA1, SART1, or BRCA1/SART1 knockdown cells (Fig. 4f, cf. DMSO vs DRB in siBRCA1-, siSART1#1-, and siBRCA1+siSART1#1-treated samples). These results indicate that SART1 promotes transcription-associated HR in an epistatic manner with BRCA1.

### The RS domain is required for SART1 recruitment to DSBs

To identify the domain or a posttranslational modification required for the recruitment of SART1 to DSBs, we prepared vectors expressing SART1 mutants. SART1 has an arginine/serine-rich domain at its amino-terminus known as the RS domain, which is shared by the SR protein family of SFs (Hertel and Graveley, 2005; Makarova et al., 2001). SART1 also possesses a leucine zipper motif that is involved in dimerization and DNA binding (Shichijo et al., 1998). Therefore, we constructed SART1 mutants that lacked the RS domain (ΔRS) or leucine zipper motif (ΔLZ) (Fig. 5a). Furthermore, SART1 has two threonine residues (T430 and T695) that are phosphorylated by ATM/ATR kinases (Matsuoka et al., 2007). Thus, we prepared two SART1 mutants that lacked (T430A/T695A) or mimicked (T430E/T695E) phosphorylation. Silent mutations were introduced into wild-type (WT) and mutant SART1 cDNAs to make them resistant to SART1 siRNA (siSART1#1). WT or mutant SART1 cDNA was then inserted into the GFP vector to visualize SART1 behavior in live cells. U2OS cells expressing mCherry-Geminin were transfected with SART1 siRNA to deplete endogenous SART1 and then transfected with the WT or mutant GFP-SART1 vector. GFP(+)/mCherry(+) cells were subjected to laser irradiation, and GFP-SART1 accumulation along the laser track was assessed. We found that the ΔRS mutant failed to accumulate along the laser track, whereas the ΔLZ, T430A/T695A, and T430E/T695E mutants showed accumulation comparable to that of the WT SART1 (Figs. 5b–c). Consistent with the results of the phosphorylation mutants, treatment with the ATM inhibitor did not influence the recruitment of the WT GFP-SART1 to the laser track (Fig. S4).

**Figure 5.**
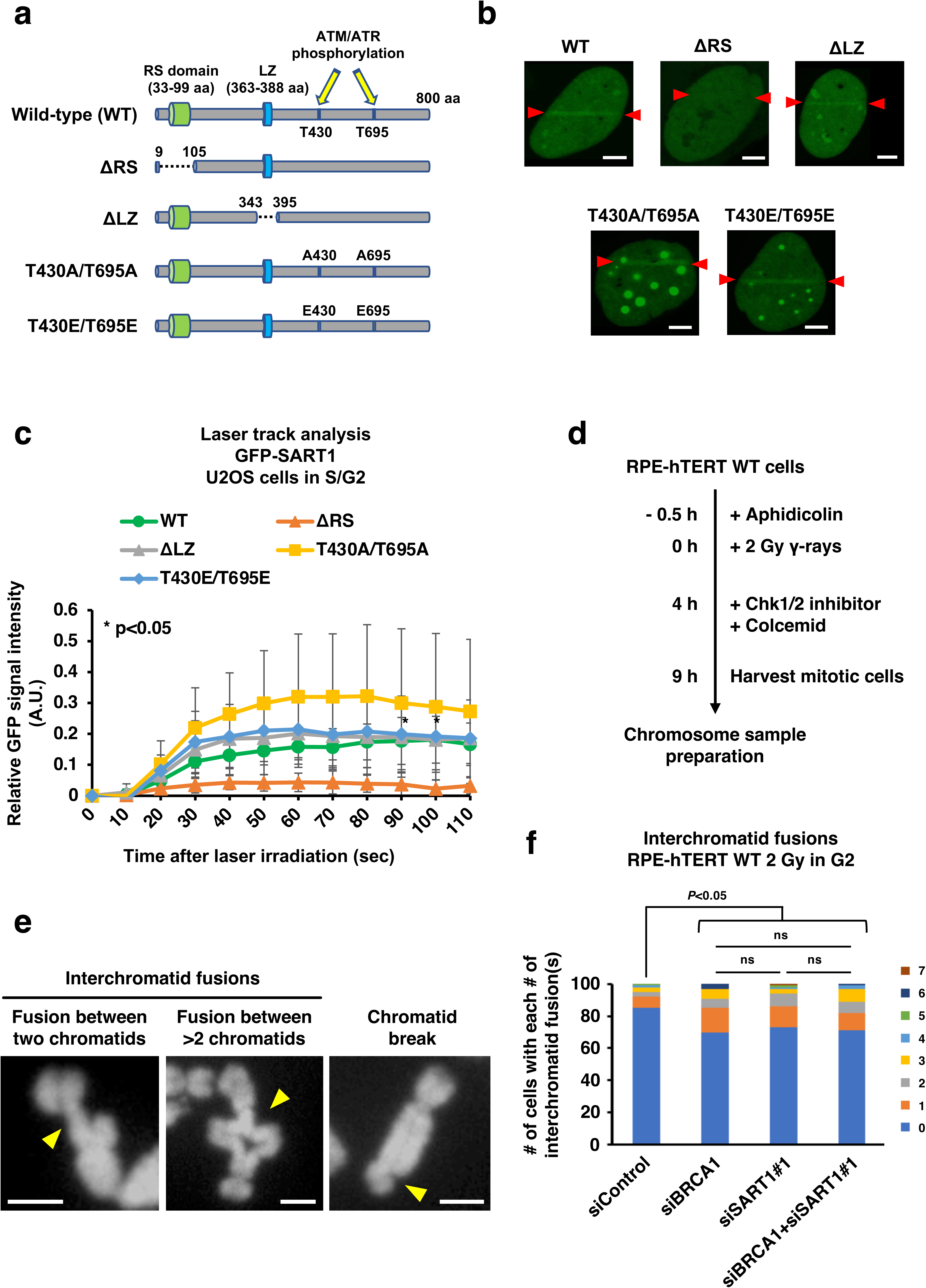
SART1 requires the RS domain for its recruitment to laser tracks and suppresses genomic alterations in an epistatic manner with BRCA1. (a) Diagram of wild-type and mutant SART1 used to test recruitment to laser tracks. The domains and ATM/ATR-dependent phosphorylation sites in SART1 are shown. (b) Live cell images of the wild-type and mutant SART1 proteins after laser irradiation. U2OS cells expressing mCherry-Geminin (an S/G2 phase marker) were transfected with wild-type or mutant GFP-SART1 vectors. One day later, GFP(+)/mCherry(+) cells were irradiated with the 730 nm laser along the line flanked by the two red arrowheads. The photosensitizer (Hoechst33342, 10 µg/mL) was added 30 m before irradiation. Scale bars: 5 µm. (c) Kinetics of recruitment of wild-type and mutant SART1 to the laser tracks. Vector transfection and laser irradiation were performed as described in (b). The intensity of GFP-SART1 in the laser-irradiated regions in S/G2-phase U2OS cells was recorded every 10 s until 110 s after irradiation. Error bars indicate the standard deviation (SD), N = 8. *, p < 0.05. (d) Scheme of chromosome experiments in (e) and (f). (e) Representative images of the detected chromosomal aberrations. Yellow arrowheads indicate aberrations. The chromosomes were stained with DAPI. Scale bars: 2 µm. (f) SART1 and BRCA1 epistatically suppress interchromatid fusion. RPE-hTERT WT cells were transfected with the indicated siRNA(s). Two days after siRNA transfection, the cells were treated, and chromosome samples were prepared as shown in (d). In total, one-hundred metaphase cells from two independent experiments (50 metaphases/experiment/sample) were analyzed for each sample. The number of aberrations in the 100 metaphases is shown.

### SART1 and BRCA1 suppress genomic alterations in the G2 phase in an epistatic manner

Finally, we examined the genomic consequences of HR defects caused by SART1 knockdown. For this purpose, mitotic chromosomes in G2-phase-irradiated cells were analyzed. To strictly analyze G2 phase-derived chromosomal aberrations, cells were treated with aphidicolin, an inhibitor of DNA synthesis, from 30 m before IR until harvest of mitotic cells to prevent S phase-irradiated cells from progressing to the G2 phase (Fig. 5d). In this analysis, two types of chromosomal aberrations were observed: interchromatid fusion and chromatid break (Fig. 5e). Interchromatid fusion and chromatid break indicate misrepair between DSBs on two or more distinct chromatids and an unrepaired DSB on a chromatid, respectively. Although the frequency of chromatid breaks was comparable among siControl-, siBRCA1-, siSART1#1-, and siBRCA1/siSART1#1-transfected cells (Table 1 and Fig. S5), single knockdown of BRCA1 and SART1, or double knockdown of these factors significantly increased the frequency of interchromatid fusion (Fig. 5f and Table 2). The frequency of interchromatid fusion was similar between BRCA1 and SART1 knockdown cells. Consistent with the epistatic relationship between BRCA1 and SART1 in HR, the double knockdown of these factors did not further increase the frequency of interchromatid fusion compared to the single knockdown of BRCA1 or SART1 (Fig. 5f). These results indicated that SART1 and BRCA1 epistatically suppressed the occurrence of genomic alterations in the G2 phase.

**Table 1.** Frequency of chromatid break in BRCA1 and/or SART1 knockdown cells. RPE-hTERT WT cells were transfected with the indicated siRNA(s). Two days after siRNA transfection, the cells were treated, and chromosome samples were prepared as shown in Fig. 5d. In total, one-hundred metaphase cells from two independent experiments (50 metaphases/experiment/sample) were analyzed for each sample. The number of cells with each number of chromatid break(s) are shown.

| # of chromatid breaks | 0 | 1 | 2 | 3 | 4 |
| --- | --- | --- | --- | --- | --- |
| siControl | 87 | 12 | 1 | 0 | 0 |
| siBRCA1 | 84 | 13 | 2 | 1 | 0 |
| siSART1#1 | 84 | 14 | 1 | 0 | 1 |
| siBRCA1+siSART#1 | 82 | 11 | 6 | 1 | 0 |

**Table 2.** Frequency of interchromatid fusion in BRCA1 and/or SART1 knockdown cells. RPE-hTERT WT cells were transfected with the indicated siRNA(s). Two days after siRNA transfection, cells were treated, and chromosome samples were prepared as shown in Fig. 5d. In total, one-hundred metaphase cells from two independent experiments (50 metaphases/experiment/sample) were analyzed for each sample. The number of cells with each number of interchromatid fusion(s) are shown.

| # of interchromatid fusion(s) | 0 | 1 | 2 | 3 | 4 | 5 | 6 | 7 |
| --- | --- | --- | --- | --- | --- | --- | --- | --- |
| siControl | 85 | 7 | 3 | 3 | 1 | 1 | 0 | 0 |
| siBRCA1 | 70 | 15 | 6 | 6 | 0 | 0 | 3 | 0 |
| siSART1#1 | 73 | 13 | 8 | 3 | 1 | 1 | 0 | 1 |
| siBRCA1+siSART1#1 | 71 | 11 | 7 | 8 | 2 | 0 | 1 | 0 |

## Discussion

In this study, we demonstrated that SART1 promotes HR by facilitating DSB end resection in an epistatic manner with BRCA1. The results showed that SART1 promotes the recruitment of the HR-promoting portion of BRCA1 to DSBs, and that SART1 and BRCA1 epistatically promote 53BP1 dephosphorylation and RIF1 release from DSBs. Laser microirradiation experiments revealed that SART1 is recruited to DSBs in a BRCA1-dependent manner, and that the RS domain is required for SART1 recruitment to DSBs. This study also provides evidence that SART1 is involved in transcription-associated HR with BRCA1. Moreover, chromosome analysis demonstrated that SART1 and BRCA1 cooperatively suppressed genomic alterations after DSB induction in the G2 phase.

Previous studies have reported that many SFs are involved in HR (Adamson et al., 2012; Anantha et al., 2013; Awwad et al., 2023; Herr et al., 2015; MacHour et al., 2021; Marchesini et al., 2017; Onyango et al., 2017; Prados-Carvajal et al., 2018; Savage et al., 2014; Tanikawa et al., 2016). While many SFs promote HR by supporting the proper expression of HR factors, some SFs seem to have a direct role in HR, in addition to their canonical function in splicing (Awwad et al., 2023; Prados-Carvajal et al., 2018). We consider that SART1 directly promotes HR because knockdown of this SF did not affect the expression of HR-related factors (Figs. S1–2). The direct role of SART1 in HR is also indicated by the rapid recruitment of this SF to laser-induced DSBs in the S/G2 phase cells (Figs. 4a–b). Previous studies have reported that several SFs, such as PRP19 and PPM1G, are also recruited to laser-induced DSBs (Beli et al., 2012; Maréchal et al., 2014). However, this phenotype is not common to all SFs because some SFs do not accumulate in laser-irradiated regions (e.g., PRP3) or are excluded from (e.g., THRAP3) laser tracks (Beli et al., 2012; Maréchal et al., 2014). Therefore, it is likely that recruitment to DSB sites is limited to SFs that are directly involved in DSB repair or response.

Previous studies have investigated the role of BRCA1 in resection and established that BRCA1 promotes resection by counteracting 53BP1/RIF1-mediated resection blockade (Scully et al., 2019; Shibata, 2017; Tarsounas and Sung, 2020). Our data demonstrated that SART1 is also involved in this process because (1) the reduced number of RPA foci in SART1 knockdown cells was restored by 53BP1 knockdown and (2) the foci numbers of phosphorylated 53BP1 and RIF1 were increased by SART1 knockdown, which were both similarly observed in BRCA1 knockdown cells (Figs. 3a, c–f). SART1 is likely to contribute to BRCA1 antagonism against 53BP1/RIF1 by recruiting the HR-promoting portion of BRCA1 to DSBs because SART1 knockdown impaired BRCA1 foci formation in Rap80-knockout cells, where only HR-promoting BRCA1 complexes are present (Figs. 2c–d) (Li and Greenberg, 2012). Moreover, the laser microirradiation experiment demonstrated that SART1 requires BRCA1 for efficient recruitment to laser-induced DSBs (Fig. 4c). Thus, our results indicate that upon DSB induction, SART1 is rapidly recruited to DSBs in a BRCA1-dependent manner, and then, SART1 recruits HR-promoting BRCA1 complexes to DSBs, thereby relieving the resection barrier posed by 53BP1 and RIF1.

We previously demonstrated that BRCA1 is involved in transcription-associated HR, in which a subset of DSBs require ongoing transcription for end resection (Yasuhara et al., 2018). Our data indicate that SART1 participates in the resection step of transcription-associated HR in an epistatic manner with BRCA1 (Fig. 4f). The involvement of SART1 in transcription-associated HR was also supported by the transcription-dependent accumulation of SART1 at laser-induced DSBs (Figs. 4d–e). Thus, when DSBs occur in transcriptionally active genomic regions, SART1 is recruited to such DSBs and recruits BRCA1, which facilitates resection by promoting 53BP1 dephosphorylation and RIF1 release from DSBs (Fig. 6).

**Figure 6.**
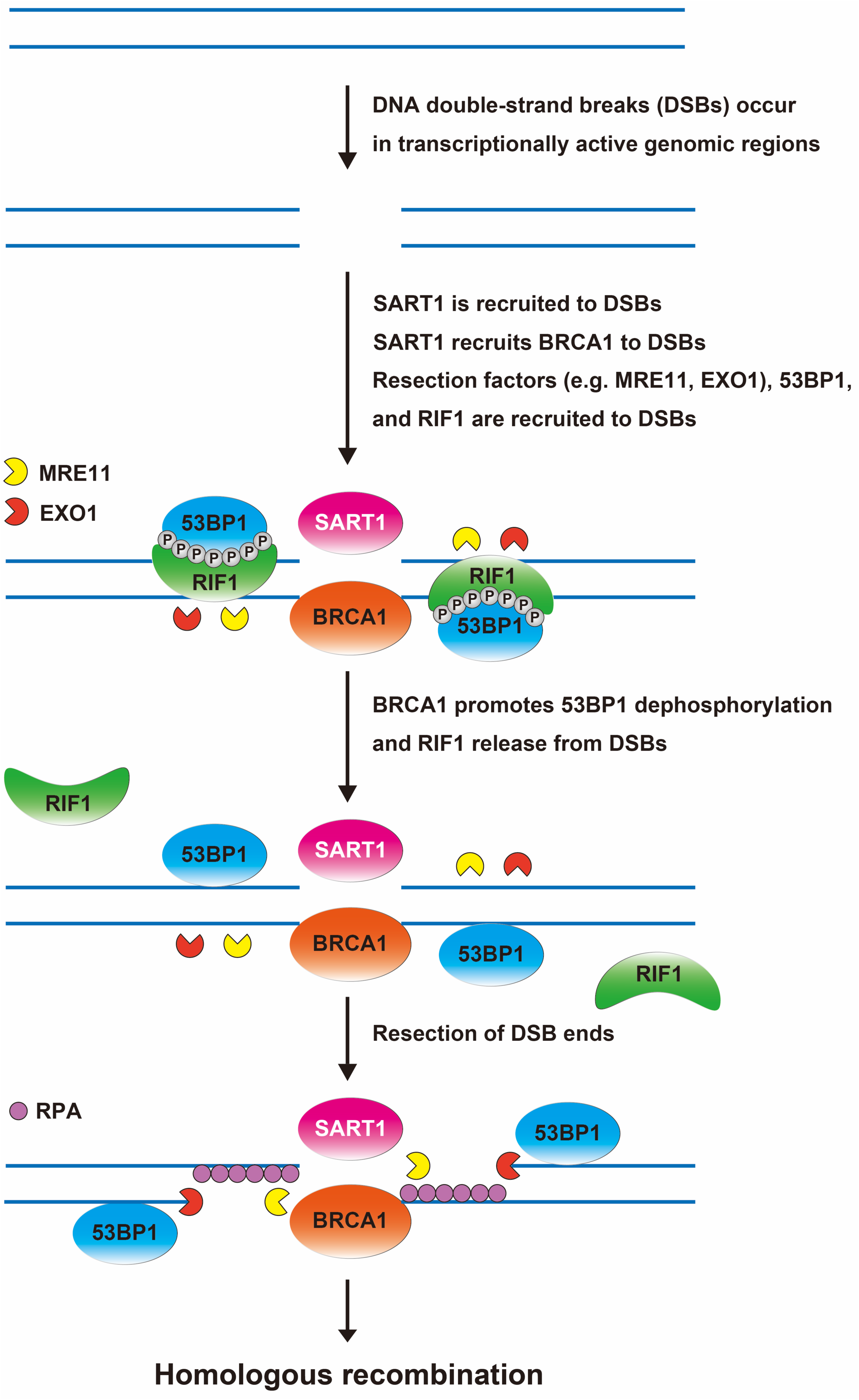
Model of the role of SART1 in promoting HR. When DSBs occur in transcriptionally active genomic regions, SART1 is rapidly recruited to DSBs in a BRCA1-dependent manner. Then, SART1 recruits BRCA1 to DSBs. Resection factors (e.g., MRE11 and EXO1), 53BP1 and RIF1 are also recruited. At the DSB sites, BRCA1 promotes 53BP1 dephosphorylation and RIF1 release from DSBs, thereby facilitating resection and subsequent HR. DNA, blue line.

Previous studies have reported that DNA breaks in BRCA1-deficient cells tend to form radial chromosomes, which is a type of interchromatid fusion (Bunting et al., 2010; Venkitaraman, 2004). Consistently, our chromosome analysis demonstrated that BRCA1 knockdown increased the frequency of interchromatid fusion after IR in the G2 phase (Fig. 5e–f and Table 2). Moreover, our results indicated that BRCA1 and SART1 epistatically suppressed this type of genomic alteration because the frequency of interchromatid fusion was similarly increased by the single knockdown of BRCA1 and SART1, and it was not further increased when both factors were knocked down (Fig. 5e–f and Table 2). Given the epistatic relationship between SART1 and BRCA1 in resection, it is likely that these factors collaboratively direct the repair pathway toward HR by promoting resection; otherwise, a more error-prone, non-HR pathway operates, which increases genome instability.

Our laser microirradiation experiments identified the RS domain as essential for SART1 recruitment to laser-induced DSBs (Fig. 5a–c). Although the mechanism by which the RS domain recruits SART1 to DSBs remains undetermined, our preliminary survey using an online predictor of the intrinsically disordered regions (IDR) showed that the N-terminal region (1-125 aa) of SART1, which includes the RS domain (33-99 aa), is highly disordered. Recent studies have revealed that proteins with an IDR tend to form globular membraneless structures called condensates (Spegg and Altmeyer, 2021). Consistently, SART1 reportedly forms condensates in the nucleus (Shin et al., 2018). In preliminary experiments, we observed that WT GFP-SART1, but not ΔRS GFP-SART1, formed condensates (data not shown). Thus, it would be interesting to speculate that the RS domain promotes SART1 condensate formation, which compartmentalizes HR factors, including BRCA1, thereby excluding 53BP1/RIF1 from DSB ends and facilitating resection and subsequent HR.

In summary, this study uncovered a mechanism by which SART1 promotes HR, which, to the best of our knowledge, has not been reported previously. Since SF genes are mutated or aberrantly expressed in various types of cancer and hematopoietic malignancies, cancer therapies targeting SFs have recently emerged. Current strategies include direct inhibition of SFs (e.g., SF3B1) or indirect inhibition through the modulation of post-translational modifications of SFs (e.g., methylation, phosphorylation, and ubiquitylation) by upstream modifiers (Bradley and Anczuków, 2023). Because many SFs are involved in DNA repair, a combination of SF inhibitors and DNA-damaging cancer therapies, such as radiotherapy and platinum-based therapies, is expected to improve the efficacy of cancer therapies. Thus, further studies on the roles of SFs in DNA repair will contribute to the development of SF inhibition-based cancer therapies.

## Materials and Methods

### Cell culture and irradiation

Human telomerase reverse transcriptase-immortalized retinal pigment epithelial cells (RPE-hTERT) were cultured in Dulbecco’s modified Eagle’s medium (DMEM)/Ham’s F12 medium supplemented with 10% fetal bovine serum (FBS). U2OS cells were cultured in α-MEM supplemented with 10% FBS. Cells were subjected to gamma-ray irradiation using a ^137^Cs source.

### siRNA transfection

For RPE-hTERT cells, siRNA transfection was performed using DharmaFECT 4 (Horizon Discovery, Cambridge, UK), following the manufacturer’s instructions. Briefly, 1 × 10^5^ cells were seeded in a 35-mm dish and cultured for 1 d. Next, siRNA and DharmaFECT 4 were separately diluted in Opti-MEM, mixed, and incubated for 20 m at room temperature (RT). Cells were incubated with a mixture of siRNA and DharmaFECT 4 diluted in culture medium containing 20% FBS for 1 d. The medium was then replaced with fresh medium containing 10% FBS, and the cells were cultured for another day until the experiment. For U2OS cells, reverse siRNA transfection was performed using Lipofectamine RNAiMAX (Thermo Fisher Scientific, MA, USA). First, siRNA and Lipofectamine RNAiMAX were separately diluted in Opti-MEM, mixed, and incubated for 20 m at RT. During the incubation, the cells were harvested by trypsinization and suspended in the medium at a concentration of 2 × 10^5^ cells/mL. The cells were then mixed with diluted siRNA/lipofectamine RNAiMAX, plated in a 35-mm dish, and cultured for 1 d. The medium was then replaced with fresh medium containing 10% FBS, and the cells were cultured for another day until the experiment. Two days after siRNA transfection, the cells were subjected to further experiments, including irradiation, fixation, and immunofluorescence. The siRNA sequences used in this study are listed in Table S1.

### DR-GFP assay

U2OS cells harboring the DR-GFP HR reporter were used to examine the efficiency of HR. The cells were transfected with the indicated siRNAs and cultured overnight. Next, the cells were transfected with the pCBASce I vector (expression vector of I-Sce I endonuclease) using Nano Juice (Merck, NJ, USA). Cells were harvested and suspended in PBS containing 1 mM EDTA 48 h after pCBASce I transfection. The percentage of GFP-positive cells was determined using an Attune NxT flow cytometer (Thermo Fisher Scientific, MA, USA).

### Immunofluorescence analysis, foci counting, and image acquisition

Cells were plated on coverslips in 35-mm dishes. The cells that had undergone DNA synthesis were labeled with 10 μM EdU, a thymidine analog, from 30 m before irradiation until fixation. For RPA foci, cells were pre-extracted with 0.2% Triton X-100 in phosphate-buffered saline (PBS) for 1 m (Figs. 1e, f, 2a, and 3a) or pre-extracted twice for 3 m with CSK buffer (10 mM HEPES, 100 mM NaCl, 300 mM sucrose, 3 mM MgCl_2_, and 0.7% Triton X-100) containing 0.3 mg/mL RNase A (Fig. 4f), followed by fixation with 3% paraformaldehyde (PFA)/2% sucrose in PBS for 10 m (Figs. 1e, f, 2a, and 3a) or 2% PFA in CSK buffer for 10 m (Fig. 4f) at RT. For RAD51 foci, the cells were pre-extracted with 0.2% Triton X-100 in PBS for 1 m, followed by fixation with 3% PFA/2% sucrose in PBS for 10 m at RT. For BRCA1 foci, cells were pre-extracted with CSK buffer for 5 m, followed by fixation with 4% PFA in PBS for 10 m at RT. To detect 53BP1 pT543 and RIF1 foci, the cells were fixed with 3% PFA/2% sucrose in PBS for 10 m at RT, followed by permeabilization with 0.2% Triton X-100 in PBS for 2.5 m. Cells were incubated with primary antibodies for 30 m at 37 °C, followed by incubation with Alexa Fluor 488- or 555-conjugated secondary antibodies for 30 m at 37 °C. The primary antibodies used for the immunofluorescence analysis are listed in Table S2. All antibodies were diluted in 2% bovine serum albumin in PBS. Next, the samples were incubated with an EdU detection solution (Alexa Fluor 647 azide in 100 mM Tris-HCl, 100 mM L-ascorbic acid, and 4 mM CuSO_4_) for 30 m at RT in the dark. Coverslips were mounted onto glass slides with Vectashield mounting medium containing 4′,6-diamidino-2-phenylindole (DAPI) (Vector Laboratories, CA, USA). The foci of RPA, RAD51, BRCA1, 53BP1 pT543, and RIF1 in G2 phase cells were counted using a fluorescence microscope (BX53, Olympus, Tokyo, Japan or BZ-X700, Keyence, Tokyo, Japan). Images of the foci were acquired using a confocal fluorescence microscope (LSM800; Zeiss, Oberkochen, Germany) with a 100× objective. Z-stack images were captured at intervals of 0.2–0.25 µm. Maximal intensity projection images were obtained using ZEN 2.3 image analysis software (Zeiss).

### Establishment of Rap80 knockout cell line

Rap80 knockout RPE-hTERT (RPE-hTERT ΔRap80) cells were established as previously described (Yasuhara et al., 2018). Briefly, the All-in-One CRISPR-Cas9D10A vector (Addgene #74119) comprising the guide RNA targeting Rap80 and GFP-labeled Cas9D10A nickase was transfected into the cells. Transfected cells were enriched using FACS Aria II (Becton Dickinson) and seeded in 96-well plates. Rap80 knockout clones were screened using polymerase chain reaction (PCR) and further confirmed by western blotting.

### Laser track analysis

One day before analysis, U2OS cells expressing mCherry-Geminin seeded on a 35-mm glass-bottom dish (Matsunami, Osaka, Japan) were transfected with the GFP-SART1 vector using Lipofectamine 2000 (Thermo Fisher Scientific, USA), following the manufacturer’s instructions. Next, the cells were incubated with 10 µg/mL Hoechst33342 (FUJIFILM Wako, Osaka, Japan) for 10 m and incubated at 37 °C in a temperature-controlled chamber (Zeiss) during the analysis. GFP(+)/mCherry(+) cells were irradiated with a Mai Tai laser (Spectra-Physics, Santa Clara, CA, USA) at a wavelength of 730 nm and a nominal power of approximately 10 mW along a track with a width of approximately 500 nm. The cells were observed under an LSM510 microscope (Zeiss) equipped with a 63× objective. For each condition, three to seven cells from two biological replicates were imaged at intervals of 10 s. For the laser track analysis of mutant GFP-SART1, U2OS cells expressing mCherry-Geminin were transfected with SART1 siRNA (siSART1#1) and cultured for 1 d to deplete endogenous SART1. The cells were then transfected with siSART1#1-resistant GFP-SART1 vector (wild-type or mutant), cultured for another day, and subjected to laser track analysis.

### Cloning of SART1 cDNA and generation of SART1 mutants

Human SART1 mRNA (NM_005146) was reverse-transcribed into complementary DNA (cDNA), which was cloned into the pEGFP-C3 vector (Clontech, Mountain View, CA, USA) using the In-Fusion HD cloning kit (Takara Bio, Shiga, Japan). SART1 mutants were generated using the KOD-Plus-Mutagenesis kit (Toyobo, Osaka, Japan) with some modifications. Mutations were confirmed by Sanger sequencing. For laser track analysis, silent mutations were introduced into wild-type and mutant GFP-SART1 cDNAs to make them resistant to SART1 siRNA (siSART1#1).

### Preparation of the chromosome samples

Chromosomal samples were prepared as previously described, with some modifications (Yamauchi et al., 2011; Yamauchi et al., 2017). Briefly, siRNA-treated RPE-hTERT cells were plated in two T75 flasks per sample. To analyze chromosomes only in G2-irradiated cells, cells were treated with aphidicolin (3 µM), a DNA synthesis inhibitor, from 30 m before irradiation until fixation, so that the irradiated S phase cells did not progress to the G2 phase. Thirty minutes after the addition of aphidicolin, cells were irradiated with 2 Gy γ-rays and 4 h later, Chk1/2 inhibitor (SB218078, 2.5 µM) and colcemid (0.025 µg/mL) were added to the culture medium. Chk1/2 inhibitor and colcemid were used to avoid bias of the G2 checkpoint and to enrich mitotic cells, respectively. Five hours later (9 h after irradiation), mitotic cells were shaken off from the flasks, suspended in medium, and subjected to hypotonic treatment for 20 m at 37 °C in 0.075 M potassium chloride, followed by fixation with ice-cold Carnoy’s fixative (methanol: acetic acid = 3:1). After centrifugation (1,200 rpm, 2 m, 4 °C), the cells were suspended in Carnoy’s fixative, dropped onto glass slides, and dried for 1–3 d. Chromosomes were stained with DAPI in the Vectashield mounting medium.

### Statistical analysis

All statistical analyses, except for the analysis of chromosomal aberration data (Fig. 5f), were performed using Prism 8 software (GraphPad Software, San Diego, CA, USA). The means between two groups were compared using unpaired t-test, Welch’s t-test, or two-tailed Mann– Whitney test, whereas those between more than two groups were compared using Dunn’s multiple comparisons test. For the analysis of the chromosomal aberration data in Fig. 5f, a zero-inflated Poisson (ZIP) regression model was applied using the pscl package in R, because the data contained excessive zeros. Differences were considered statistically significant at P < 0.05.

## Acknowledgments

We thank Dr. Yasuyoshi Oka for providing information on the anti-SART1 antibody. We thank the Research Promotion Unit of the Medical Institute of Bioregulation at Kyushu University for technical assistance. We appreciate the technical assistance from the Research Support Center, Research Center for Human Disease Modeling, Kyushu University Graduate School of Medical Sciences, which is partially supported by the Mitsuaki Shiraishi Fund for Basic Medical Research. This work was supported by Grants-in-Aid for Scientific Research (KAKENHI) from the Japan Society for the Promotion of Science (JP18K11645 and JP22K12376 to M.Y., JP17H04713 to A.S., and JP18K18191 to T.Y.), Takeda Science Foundation, and Network-type Joint Usage/Research Center for Radiation Disaster Medical Science.

## Author contributions

M.Y. conceived the study and designed the experiments with T.Y., A.S., K.M., K.S., and N.M. R.K. and T.Y. generated RPE-hTERT ΔRap80 cells and performed laser track analysis. Y.U., R. K-I, and A.S. performed the DR-GFP assay, RNA-seq, and bioinformatic analyses. M.Y., M.H., and A.S. constructed vectors. M.Y., M.H., K.O., A. Shakayeva, and P.K. performed immunofluorescence, microscopy, and western blotting analyses. H.S. performed γ-irradiation. M.Y. and Y.A performed the chromosome experiments. M.Y. summarized all data for the figures and prepared the manuscript with support from T.Y., A.S., K.M., K.S., and N.M. M.Y. supervised this study.

## Data availability statement

The RNA-seq data have been deposited at DNA Data Bank of Japan (DDBJ) under accession number: DRA017000.

