## Supplementary materials for "Involvement of the splicing factor SART1 in the BRCA1-dependent homologous recombination repair of DNA double-strand breaks"

Supplementary Methods

Supplementary Figures and Tables (Figure S1–5, Table S1–2)

Supplementary Figure legends

#### Supplementary Methods

##### *RNA-seq*

Total RNAs from cultured RPE-hTERT cells was extracted using NucleoSpin RNA (MACHEREY-NAGEL) according to the manufacturer's protocol. Purified RNA quality was evaluated by the RNA integrity number (RIN) using the Agilent RNA6000 Pico Kit and Agilent 2100 Bioanalyzer (Agilent Technologies, Santa Clara, CA, USA). High-quality RNA samples (RNA integrity number (RIN) > 9.8) were used for RNA-seq analysis. One microgram of total RNA was used to prepare sequence libraries using a KAPA mRNA HyperPrep Kit (Kapa Biosystems Inc., Wilmington, MA, USA), following the manufacturer's instructions. The generated libraries and 1% PhiX spike-in were then subjected to paired-end sequencing of 76-bp reads using a NextSeq500 System (Illumina Inc., San Diego, CA, USA) with a NextSeq500 High Output v2.5 Kit (Illumina). Alignment of reads (approximately 43 million reads) to the GRCh38 genome using STAR (version 2.7.6a) was followed by quantification of gene and transcript levels using RSEM (version 1.3.3) based on Ensembl release 100 (Dobin et al., 2013; Li and Dewey, 2011). For the analysis, read counts > 0 were used as thresholds. Count normalization and identification of differentially expressed genes (DEGs) were conducted using the iDEGES/edgeR-edgeR pipeline in the R statistical package TCC (Sun et al., 2013). DEGs were selected based on a q-value of <0.05.

##### *Reagent*

An ATM inhibitor (KU55933; Cat. No. 118502) was obtained from Merck Millipore and treated at a final concentration of 10 μM 1 h before laser irradiation.

### Supplementary Figure S1

**a**

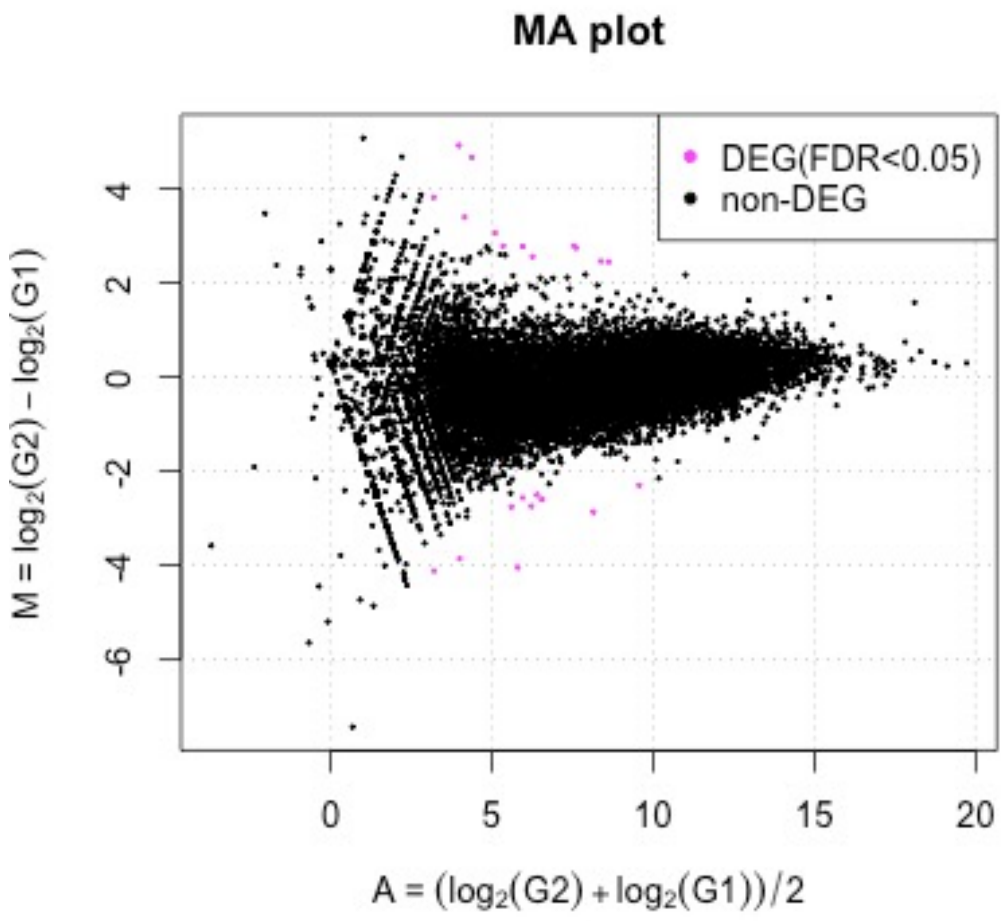


**b**

| |
|---|
| FP236241.2 |
| CU634019.2 |
| AC019117.4 |
| CU633906.2 |
| AC009412.1 |
| AL137782.1 |
| CXCL8 |
| AC034102.2 |
| GDNF |
| SLC37A2 |
| EDN1 |
| NOG |
| Z95118.2 |
| C5AR1 |
| AC087190.3 |
| SULT1A4 |
| ADAM32 |
| AC116366.3 |
| SART1 |
| GLI1 |
| SDCBP2-AS1 |
| HMGN1P3 |

### Supplementary Figure S2

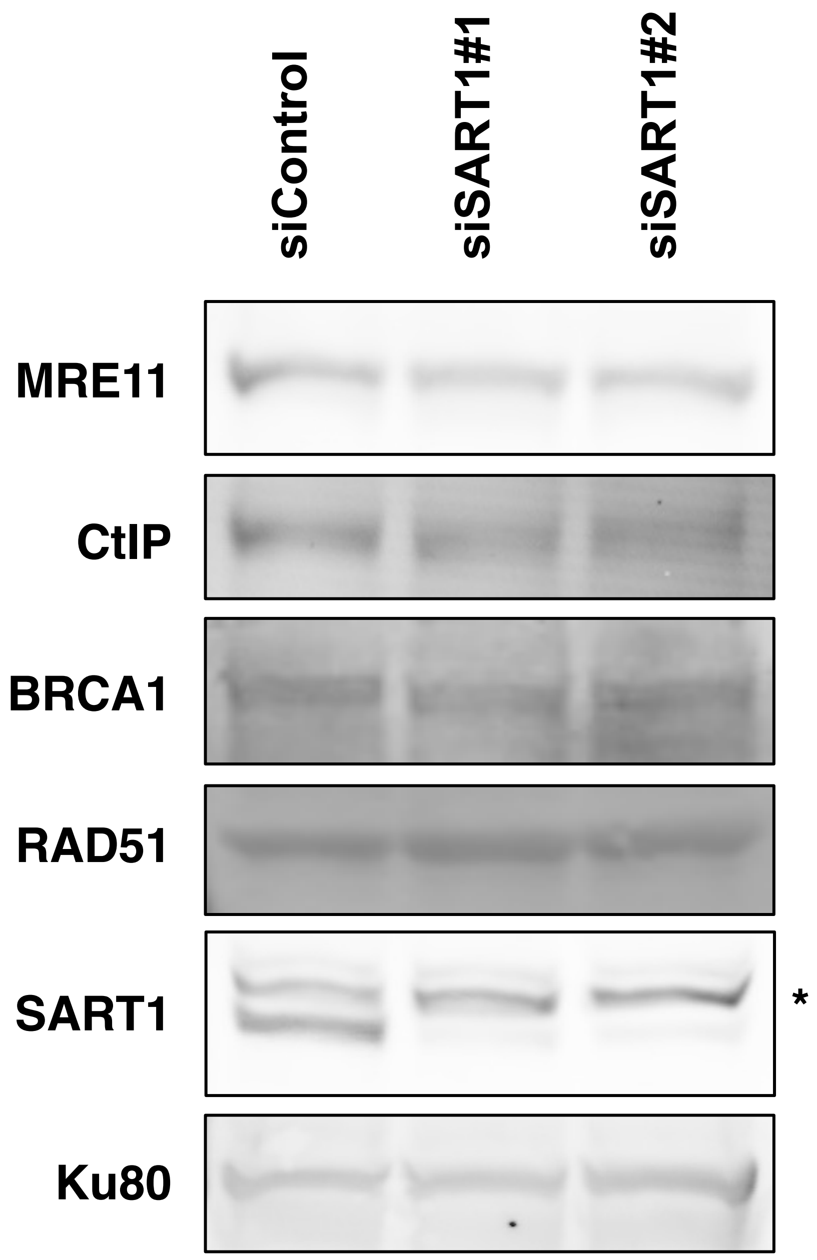


* Non-specific bands

### Supplementary Figure S3

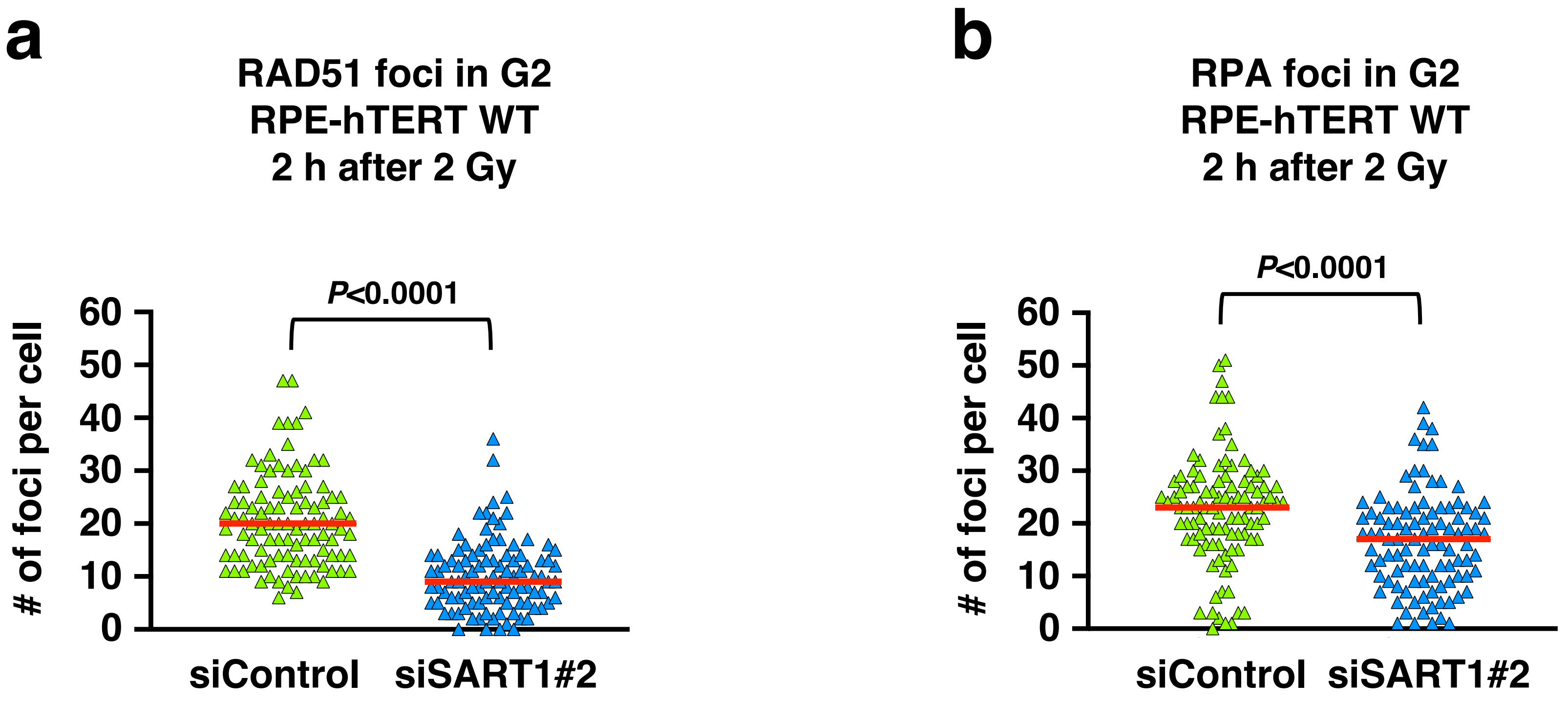

### Supplementary Figure S4

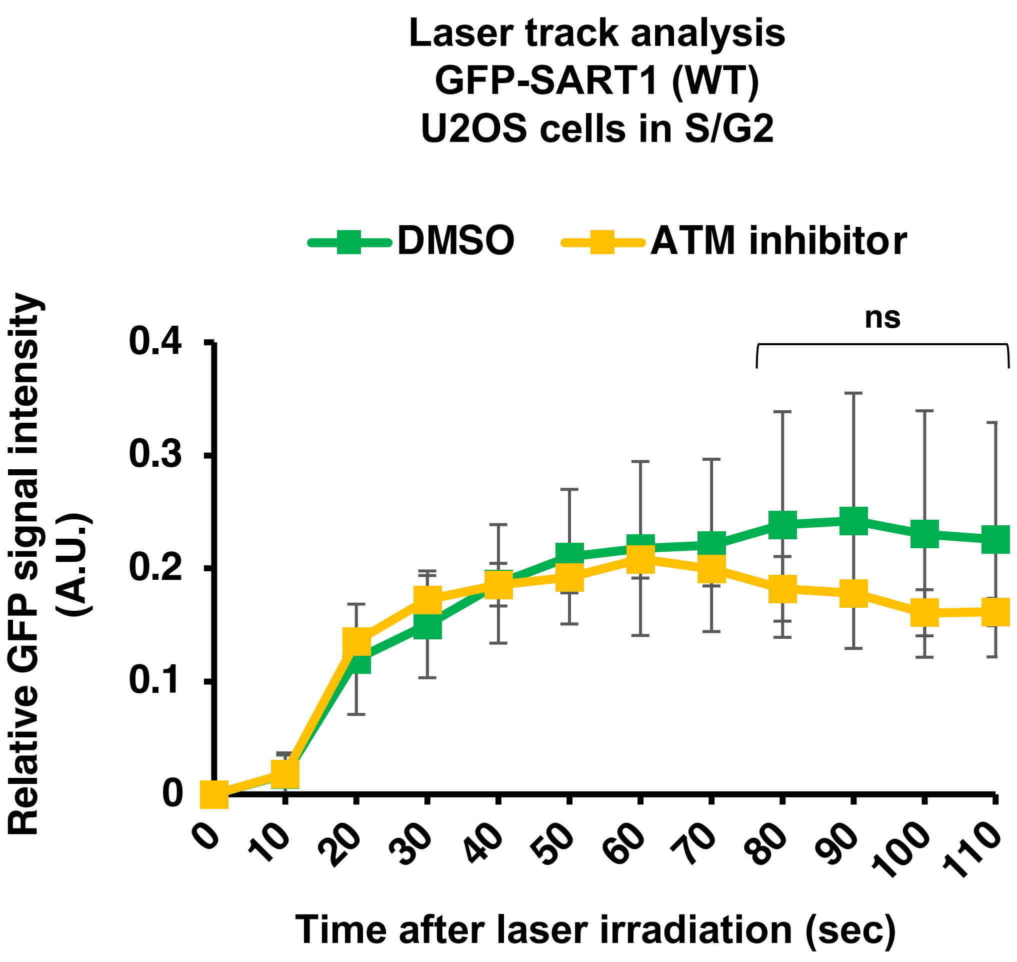

### Supplementary Figure S5

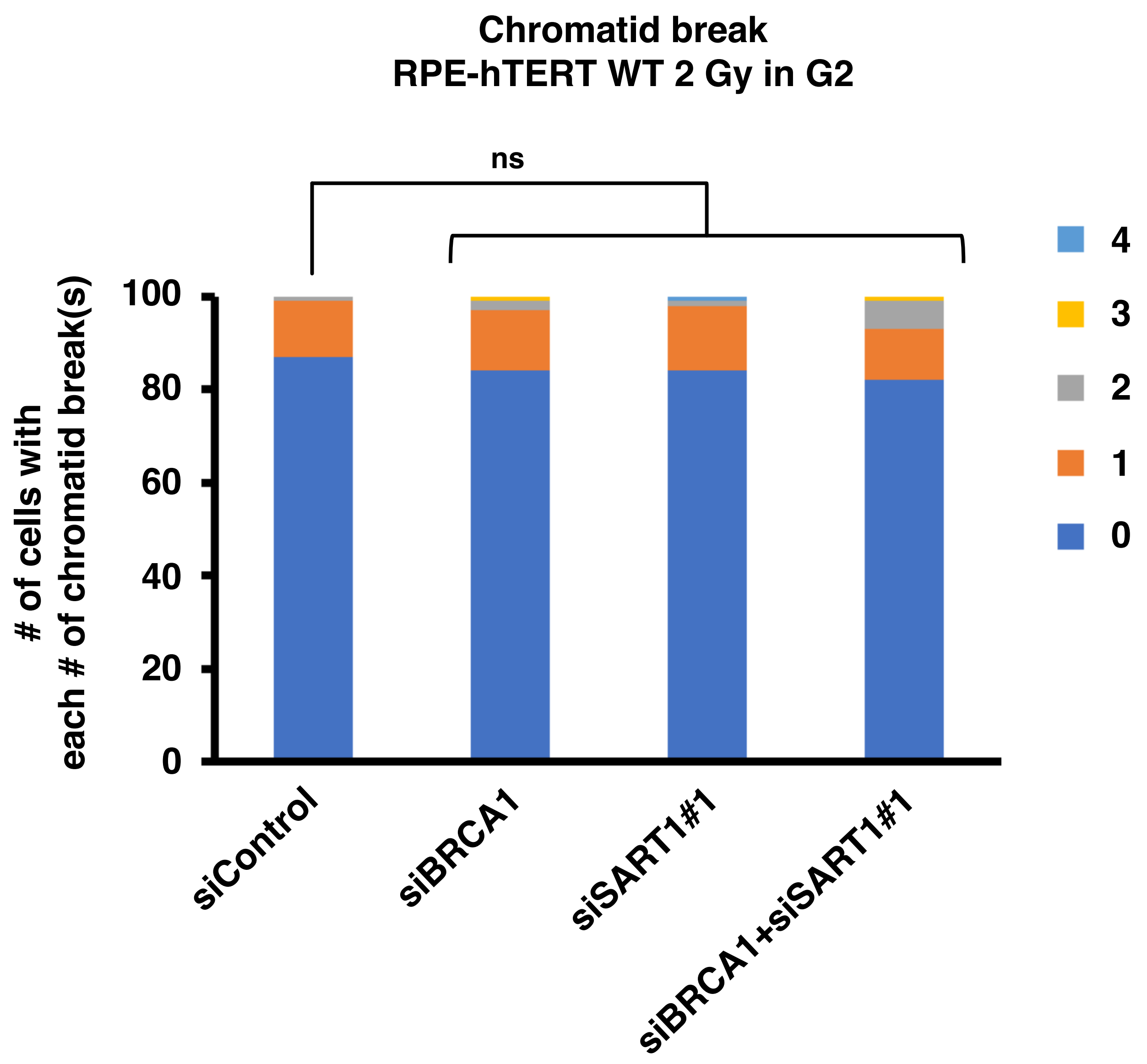

### Supplementary Table S1

**Sequence of siRNAs used in the present study**

| siRNA | Sequence of sense strand of siRNA duplex (5'-3') |
|---|---|
| siSART1#1 | CCGAAUACCUCACGCCUGAdTdT |
| siSART1#2 | GCAAGAGCAUGAACGCGAAdTdT |
| siBRCA1 | GGAACCUGUCUCCACAAAGdTdT |
| si53BP1 | GGACUCCAGUGUUGUCAUUUU |

### Supplementary Table S2

**Antibodies used in the present study**

| Target | Clone or Cat# | Host | Source | Application (dilution ratio) |
|---|---|---|---|---|
| RPA2 | LS-C38952 | Rat | Life Span BioSciences, Inc. | IF (1:500) |
| RAD51 | 14B4/GTX70230 | Mouse | GeneTex | IF (1:500)<br>WB (1:200) |
| RAD51 | ab133534 | Rabbit | Abcam | IF (1:1000) |
| BRCA1 | D-9 | Mouse | Santa Cruz Biotech. | IF (1:250) |
| BRCA1 | 07-434 | Rabbit | Merck | WB (1:200) |
| 53BP1 pT543 | 3428 | Rabbit | Cell Signaling Tech. | IF (1:500) |
| RIF1 | A300-569A | Rabbit | Bethyl Laboratories | IF (1:500) |
| CENPF | 58982 | Rabbit | Cell Signaling Tech. | IF (1:1000) |
| CENPF | 610768 | Mouse | BD Biosciences | IF (1:1000) |
| MRE11 | 31H4/4847 | Rabbit | Cell Signaling Tech. | WB (1:200) |
| CtIP | D76F7/9201 | Rabbit | Cell Signaling Tech. | WB (1:200) |
| SART1 | HPA031188 | Rabbit | Atlas antibodies | WB (1:200) |
| Ku80 | C48E7/2180 | Rabbit | Cell Signaling Tech. | WB (1:200) |

IF, Immunofluorescence; WB, Western blotting

**Supplementary Figure legends**

Figure S1. Transcriptome analysis of SART1 knockdown cells using RNA-seq.

(a) MA plot comparing the transcriptome of SART1 knockdown cells and control cells. Data were obtained using RNA-seq and analyzed as described in the Supplementary Methods.

(b) Genes that were differentially expressed in SART1 knockdown cells compared with control cells.

Figure S2. Effect of SART1 knockdown on protein levels of major HR factors.

RPE-hTERT cells were transfected with indicated siRNAs. Two days later, cells were lysed with 2x Laemli sample buffer (Merck, NJ, USA). The levels of the indicated proteins were examined by western blotting.

Figure S3. SART1 promotes HR and resection of DSBs in the G2 phase.

(a) Number of RAD51 foci in SART1 knockdown cells. Wild-type RPE-hTERT cells (RPE-hTERT WT) were transfected with the indicated siRNA. Two days after siRNA transfection, the cells were irradiated with 2 Gy γ-rays and fixed 2 h later. The cells were treated with EdU from 30 m before irradiation until fixation to label S phase cells. The fixed cells were subjected to RAD51/CENPF immunofluorescence and EdU detection.

(b) Number of RPA foci in SART1 knockdown cells. The siRNA transfection, EdU treatment, irradiation, and fixation were performed as described in (a). The fixed cells were subjected to RPA/CENPF immunofluorescence and EdU detection.

In (a) and (b), the number of foci in 100 G2 cells from two independent experiments (50 G2 cells/experiment/sample) is shown. Each symbol in (a) and (b) represents the number of foci per cell. Red bars in (a) and (b) indicate the median number of foci in each sample.

Figure S4. Effect of ATM inhibition on the recruitment of SART1 to the laser track.

U2OS cells expressing mCherry-Geminin (an S/G2 phase marker) were transfected with the wild-type GFP-SART1 vector. One day later, GFP(+)/mCherry(+) cells were irradiated with the 730 nm laser. The ATM inhibitor (KU55933, 10 μM) and the photosensitizer (Hoechst33342, 10 μg/mL) were added 30 m before irradiation. The intensity of GFP-SART1 in the laser-irradiated regions of S/G2-phase U2OS cells was recorded every 10 s until 110 s after irradiation.

Figure S5. Frequency of chromatid breaks in BRCA1 and/or SART1 knockdown cells.

RPE-hTERT WT cells were transfected with the indicated siRNA(s). Two days after siRNA transfection, the cells were treated and chromosome samples were prepared, as shown in Fig. 5d. In total, one-hundred metaphase cells from two independent experiments (50 metaphases/experiment/sample) were analyzed for each sample. The number of aberrations in the 100 metaphases is shown.
